# An improved experimental pipeline for preparing circular ssDNA viruses for next-generation sequencing

**DOI:** 10.1101/2020.09.30.321224

**Authors:** Catherine D. Aimone, J. Steen Hoyer, Anna E. Dye, David O. Deppong, Siobain Duffy, Ignazio Carbone, Linda Hanley-Bowdoin

## Abstract

We present an optimized protocol for enhanced amplification and enrichment of viral DNA for Next Generation Sequencing of begomovirus genomes. The rapid ability of these viruses to evolve threatens many crops and underscores the importance of using next generation sequencing efficiently to detect and understand the diversity of these viruses. We combined enhanced rolling circle amplification (RCA) with EquiPhi29 polymerase and size selection to generate a cost-effective, short-read sequencing method. This optimized protocol produced short-read sequencing with at least 50% of the reads mapping to the viral reference genome. We provide other insights into common misconceptions about RCA and lessons we have learned from sequencing single-stranded DNA viruses. Our protocol can be used to examine viral DNA as it moves through the entire pathosystem from host to vector, providing valuable information for viral DNA population studies, and would likely work well with other CRESS DNA viruses.

**Highlights:**

- Protocol for short-read, high throughput sequencing of single-stranded DNA viruses using random primers
- Comparison of the sequencing of total DNA versus size-selected DNA
- Comparison of phi29 and Equiphi29 DNA polymerases for rolling circle amplification of viral single-stranded DNA genomes

## 1. Introduction

*Begomoviruses*, one of the nine genera in the *Geminiviridae*, are single-stranded DNA (ssDNA) viruses that infect a wide variety of plant species, including many important crops. They are also classified as CRESS DNA viruses, a large group of circular ssDNA viruses that encode replication-associated proteins (Rep) originating from a common ancestor (Zhao et al., 2019). Eukaryotic CRESS DNA viruses impact a wide range of plant and animal hosts and evolve rapidly (Zhao et al., 2019).

Cassava and tomato are among the important crops whose yields are severely impacted by begomovirus diseases. Cassava is an important root crop in Africa, Asia, and Latin America, with African farmers producing over half of the total cassava worldwide (FAOSTAT, 2016). In Africa and more recently in Asia, cassava yields have been reduced by Cassava mosaic disease (CMD), which is caused by a complex of 11 begomoviruses collectively referred to as cassava mosaic begomoviruses (CMBs). Annual cassava losses in Africa have been estimated to be 15-24% or 12-23 million tons (US $1.2-2.3 billion) (Thresh J. M., 1997; Uzokwe et al., 2016). In some regions of Africa, cassava farmers have experienced losses of up to 95%. Tomato, an important vegetable crop that is grown around the world, is a host for over 100 begomovirus species, with the most devastating being tomato yellow leaf curl virus (Moriones and Navas-Castillo, 2000). In the eastern United States, tomato production is also negatively impacted by a second begomovirus, tomato mottle virus (ToMoV), which causes widespread disease with yield losses of up to 50% (Abouzid, Polston, and Hiebert, 1992; Polston and Anderson, 1997). Given their significant impact on agriculture, it is important to understand how begomoviruses change over time and adapt to new hosts and environments. Next generation sequencing (Jeske, 2018) is an important approach for gaining insight into begomovirus populations.

Begomoviruses fall into two classes – the Old World viruses and the New World viruses (Lefeuvre et al., 2011). Their genomes consist of either one or two circular DNAs. CMBs are Old World viruses, while ToMoV is a New World virus. The genomes of the CMBs and ToMoV consist of two components designated as DNA-A and DNA-B that together total 5-6 Kb in size. Both components are required for systemic infection (Stanley and Gay, 1983). The genome components contain divergent transcription units separated by a 5’ intergenic sequence that contains the origin of replication and promoters for gene transcription (Hanley-Bowdoin et al., 2013). DNA-A encodes 5-6 proteins necessary for replication, transcription, encapsidation, and combatting host defenses (Hanley-Bowdoin et al., 2013). DNA-B encodes two proteins essential for movement (Hanley-Bowdoin et al., 2013).

Begomoviruses are encapsidated into double icosahedral virions and transmitted by whiteflies (*Bemisia tabaci*) (Hanley-Bowdoin et al., 2013). When a whitefly feeds on the phloem of an infected plant, it acquires virions that can be transmitted to a healthy plant during the next feeding cycle. Structural studies have shown that a begomovirus virion only contains one ssDNA molecule, such that the DNA-A and DNA-B components of bipartite viruses are packaged separately into virions (Bottcher et al., 2004). As a consequence, successful transmission of a bipartite begomovirus requires acquisition and transmission of at least two virions – one containing DNA-A and another containing DNA-B. Once the virions enter a phloem-associated cell, viral ssDNA is released, converted to double-stranded DNA (dsDNA), and replicated via a rolling circle mechanism (Hanley-Bowdoin et al., 2013). As infection proceeds, nascent viral ssDNA can undergo multiple rounds of replication or be packaged into virions for future transmission by whiteflies. ToMoV is only transmitted by whiteflies, while CMBs can be transmitted via vegetative propagation of infected stem cuttings as well as by whiteflies (Legg et al., 2014).

Begomoviruses have been shown to evolve rapidly (Duffy and Holmes, 2009; Lima et al., 2017; Rocha et al., 2013), making them good models for studying the evolution of ssDNA viruses. Many factors can contribute to begomovirus evolution, including agricultural practices, whitefly transmission, and abiotic stress. This underscores the importance of understanding how begomoviruses evolve through an entire pathosystem from host to vector. Generally, ssDNA viruses exist as genetically diverse populations, with variation similar to that of RNA virus populations (Elena and Sanjuán, 2007; Safari and Roossinck, 2014). The high genetic diversity of virus populations is linked to their rapid ability to evolve, to emerge in a new host, and to break disease resistance (Duffy, 2008). Viral diversity is driven by high mutation and recombination rates (Lefeuvre and Moriones, 2015; Sanjuán et al., 2010). The amount and type of genetic variation within a viral population is a direct measure of evolvability and pathogenesis (de la Iglesia and Elena, 2007; Elena, Fraile, and GarcÍa-Arenal, 2014; Elena and Sanjuán, 2007). However, our ability to study viral evolution in real-time has been limited by being able to accurately describe the genetic structure of viral populations over time (Acevedo, Brodsky, and Andino, 2014).

Deep sequencing technologies have advanced our understanding of the genetic variation of evolving virus populations, beyond virus identification and characterization of viral species (Acevedo et al., 2014). Next generation sequencing can provide insight into how viral populations, as quasi-species, are highly variable with a range of beneficial or neutral mutations that occur at low frequency (Dean et al.; Dickins and Nekrutenko, 2009). With NGS, we can actively track naturally occurring viral variants through infection, adaptation to new hosts, and host range expansion (Ruark-Seward et al., 2020). Yet, the ability of NGS to track viral variants is restricted by several technical challenges, including biased amplification, errors introduced during amplification and sequencing, and low viral read depth.

Current methods of viral amplification rely on the polymerase chain reaction (PCR) using virus-specific primers and Taq polymerase or rolling circle amplification (RCA) using random hexamers and phi29 DNA polymerase (Dean et al., 2001). Unlike RCA, sequence-specific PCR can introduce sequence bias that masks viral diversity in a population (Sipos et al., 2010). RCA amplifies circular episomes like begomovirus genomes more efficiently than linear DNA, thereby enriching for begomovirus sequences (Idris et al., 2014). RCA is less susceptible to sequence bias, less error-prone than traditional PCR, and does not fix errors in the sample to be sequenced (Lou et al., 2013; Wang et al., 2014). However, RCA produces hyper-branched, concatenated products (Lasken and Stockwell, 2007), that must be linearized by restriction enzyme digestion or mechanical shearing before NGS sequencing (Inoue-Nagata et al., 2004). Short-read sequencing in combination with RCA and size selection has improved viral read depth for RNA viruses (Acevedo and Andino, 2014; Acevedo et al., 2014). For DNA viruses, long-read sequencing of size-enriched viral DNA has been successful in reducing sequencing error (Mehta et al., 2019).

Short-read sequencing has been used in combination with RCA to enrich for CMB viral sequences in cassava (Kathurima, 2016) and *Nicotiana benthamiana* (Chen, Khatabi, and Fondong, 2019). Other begomoviruses species, including tomato leaf, curl New Delhi virus (Juárez et al., 2019) and euphorbia yellow mosaic virus (Richter et al., 2016) have also been amplified by RCA for short-read sequencing. Likewise, RCA has been used to improve read depth of mastreviruses, which constitute another genus in the *Geminiviridae* (Claverie et al., 2019). Depending on the research question, the methods cited above can return sufficient viral read depth for identifying new viruses and determining the prominent viruses in an infected plant. However, to reliably detect subconsensus viral variants in a population, a higher level of coverage is required (Juárez et al., 2019).

Here, we describe an experimental pipeline for analyzing begomovirus DNA population dynamics across a complete pathosystem constituted by the plant host and the insect vector. The pipeline combines size selection and linear amplification by an improved phi29 DNA polymerase to increase viral read coverage for diversity studies (Fig. 1).

**Fig 1.**
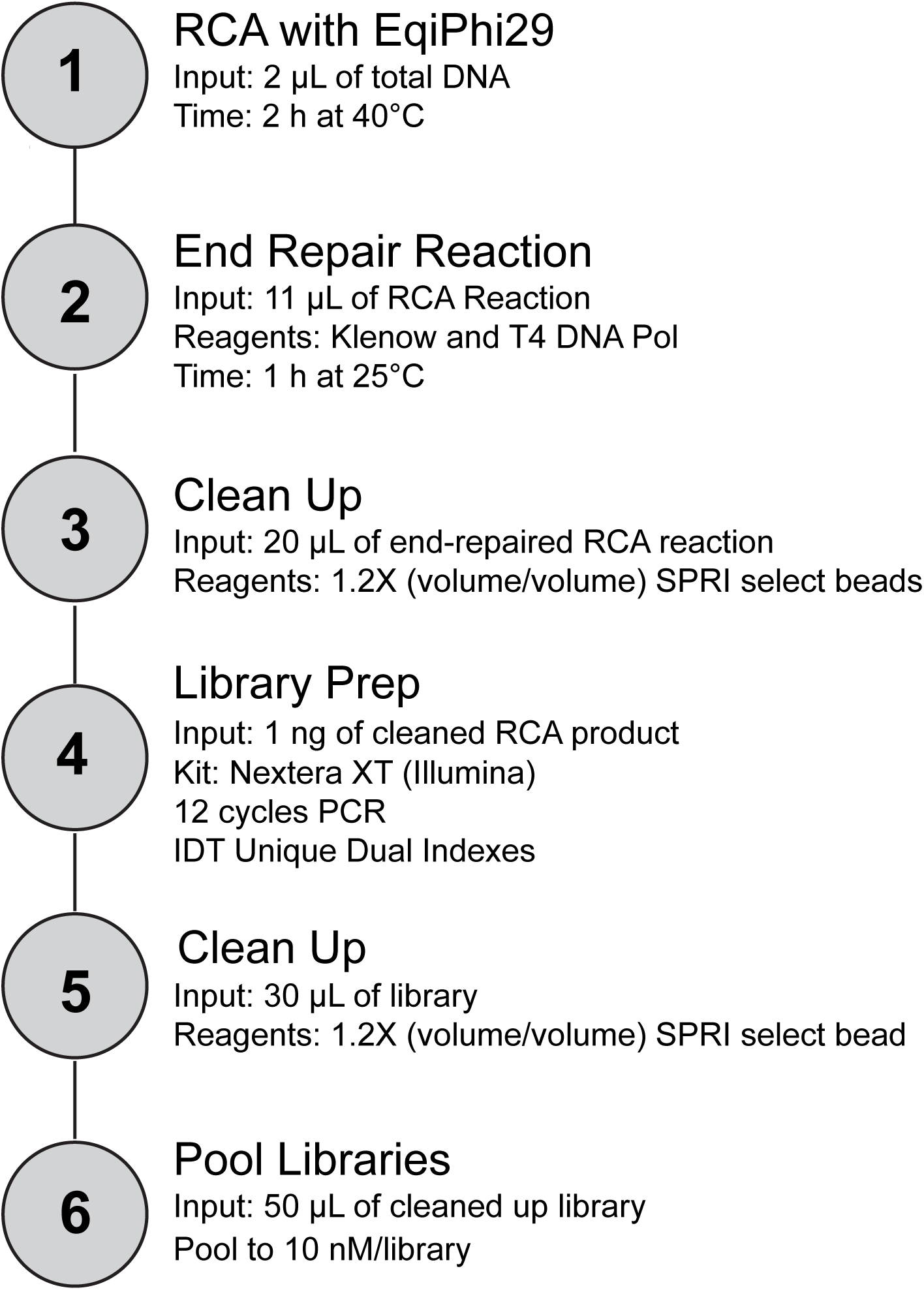
Workflow of viral DNA sequencing for short-read sequencing platforms

## 2. Methods

## 2.1 Virus-infected plants and viruliferous whiteflies

Cassava plants (*Manihot esculenta* cv. Kibandameno or Kibaha) were propagated from stem cuttings and grown at 28°C under a 12-h light/dark cycle. Plants with ca. 8-10 nodes and stems 1.5 cm in diameter (ca. 2 months after propagation) were inoculated at the apical meristem using a hand-held micro sprayer (40 psi) to deliver gold particles coated with plasmid DNA (100 ng/plasmid/plant) (Ariyo et al., 2006; Cabrera-Ponce et al., 1997). The plasmids, which contained partial tandem dimers of DNA-A and or DNA-B of *African cassava mosaic virus* (ACMV; GenBank accessions MT858793.1 and MT858794.1) and *East African cassava mosaic Cameroon virus* (EACMCV; AF112354.1 and FJ826890.1)(Chowda Reddy et al., 2012; Fondong and Chen, 2011; Fondong et al., 2000; Hoyer et al., 2020). Three plants were co-inoculated with both ACMV and EACMCV. Leaf punches from symptomatic cassava plants were sampled at 28 days post-infection (dpi), flash-frozen in liquid nitrogen, and stored for analysis.

Tomato seedlings (*Solanum lycopersicum* cv. Florida Lanai) were grown from seed at 25°C under a 12-h light/dark cycle. Plants with five true leaves (ca. 4 weeks old) were agroinoculated with ToMoV DNA-A and DNA-B (Abouzid et al., 1992; Reyes et al., 2013) as described by Rajabu et al. (2018). An infected plant was sampled at the third leaf below the apical meristem at 21 dpi, immediately prior to the whitefly access period for the acquisition of ToMoV. The leaf tissue (1 mg) was separated into two parts, one part for total DNA extraction and the other part for virion extraction. *Bemisia tabaci* MEAM1 adult whiteflies between 2 and 10 days post-eclosion were allowed to acquire the virus by feeding on a symptomatic plant infected with ToMoV for an Inoculation Access Period (IAP) of 72 h (Ng et al., 2011; Rajabu et al., 2018). Whiteflies were collected via aspiration and stored in 70% ethanol for analysis.

### 2.2 DNA extraction and size selection

Frozen leaf tissue was ground using a homogenizer (Model# MM 301, RETSCH-Laboratory Mills, Clifton, NJ), and total DNA was extracted from leaf samples using the MagMax™ Plant DNA Isolation Kit according to manufacturer’s instructions (Thermo Fisher Scientific, Waltham, MA). Total DNA was extracted from groups of five whiteflies using the Qiagen DNeasy Blood and Tissue kit according to the manufacturer’s instructions (Qiagen, Hilden, Germany). Total DNA from cassava, tomato, and whiteflies (250 ng) was size selected for 1-6 Kb DNA on a 0.75% agarose gel at 25V DC for 3-8 h using the Blue Pippin Prep system (Model # BDQ3010, Sage Science, Beverly MA). The amount of size-selected output DNA was typically less than 1 ng.

Virion DNA was generated by homogenizing five whiteflies or resuspending 1 mg of frozen cassava or tomato leaf tissue in 50 mM Tris, 10 mM MgSO4, 0.1 M NaCl, pH 7.5, followed by low-speed centrifugation. The supernatant was subjected to 0.22 μM filtration followed by DNase I digestion (2.5 U for 3 h at 37°C). Virion DNA was isolated using the QIAamp MinElute Virus Spin Kit (Qiagen, Hilden, Germany)(Ndunguru et al., 2016; Ng et al., 2011; Rosario et al., 2015).

### 2.3 Viral levels

The concentration of ACMV DNA-A (primer pair - P3P-AA2F and P3P-AA2R+4R; Table 1), DNA-B (primer pair - ACMVBdiv4 and ACMVBfor1; Table 1), and EACMCV DNA-A (primer pair – EACMVQ1 and EACMVQ; Table 1) and DNA-B (primer pair – EACMVBREV4 and EACMVBfor1.2; Table 1) were measured by quantitative PCR (qPCR) in total DNA samples (0.01 μg) extracted from cassava leaf tissue and analyzed in 96-well plates on a Max3000P System (Stratagene, San Diego CA). Primers were tested in conventional PCR to optimize annealing temperature and amplification efficiency for qPCR. For ACMV DNA-A, qPCR was performed using Power SYBR Green PCR Master Mix (Applied Biosystems, Foster City CA), starting with a 2 min denaturing step at 94°C, followed by 30 cycles consisting of 15 sec at 94°C, 1 min at 60°C, 30 sec at 72°C. The PCR conditions for EACMCV DNA-A were 10 min at 95°C, followed by 30 cycles of 30 sec at 95°C, 30 sec at 60°C, and 30 sec at 72°C. Reactions were performed in three technical replicates. ACMV DNA-B and EACMCV DNA-B were run following the above conditions respectively with an annealing temperature of 58°C. Viral DNA was quantified using a qPCR standard curve generated by amplification of a 10-fold dilution series (10^−10^ to 10^−16^ g/μL) of plasmid DNA with a single copy of ACMV DNA-A or EACMCV DNA-A following the protocol described by Rajabu et al. (2018). The concentration of the template DNA in the reaction mix was converted from ng/μL to copy number/μL using the following formula; (C × 10^−9^/MW) × NA where C = template concentration ng/μL, MW = template molecular weight in Daltons, and NA = Avogadro’s constant 6.022 × 10^23^. MW was obtained by multiplying the number of base pairs of a plasmid by the average molecular mass of one base pair (660 g/mol). A base 10 logarithmic graph of copy number versus the threshold cycle (Li et al., 2009) for the dilution factor was plotted and used as a standard curve to determine the amount of viral DNA (copy number/μL) of total DNA in a reaction mix (Rajabu et al., 2018).

**Table 1.**
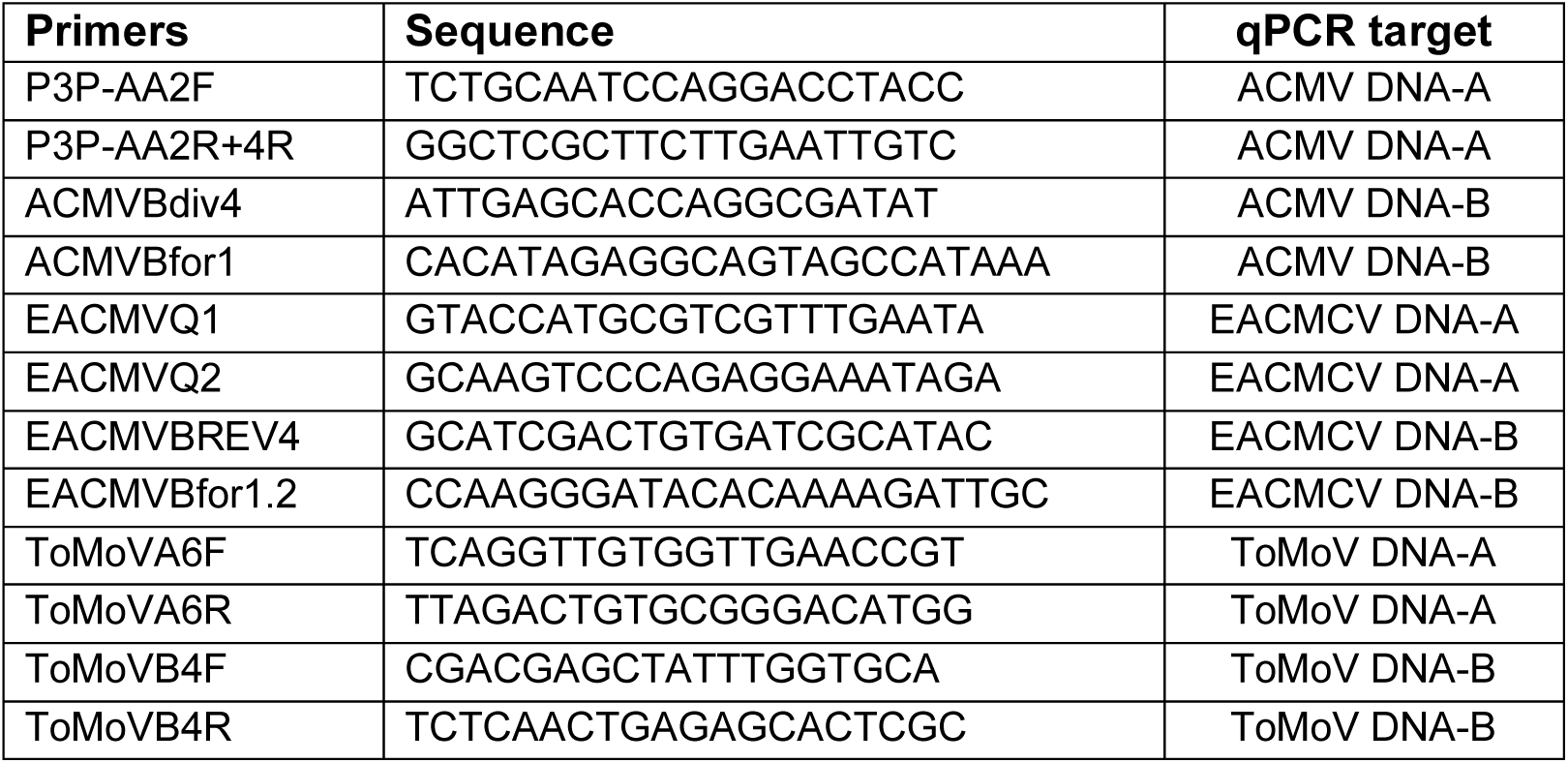
qPCR Primer Sequences.

The concentration of ToMoV genomic components was quantified by qPCR in total DNA samples (0.01 μg) from tomato leaf tissue and in whiteflies (2 ng) using the DNA-A primer pair, ToMoVA6-F, and ToMoVA6-R, and the DNA-B primer pair, ToMoVB4-F, and ToMoVB4-R (Table 1). The qPCR protocol was the same as described above for ACMV with an annealing temperature of 57°C for 1 min. Viral DNA was quantified using a qPCR standard curve generated by amplification of plasmid DNA with a single copy of each ToMoV segment (pNSB1691 and pNSB1692) following the method described above.

### 2.4 Rolling circle amplification

Total DNA (100 ng) from symptomatic Kibaha leaf tissue was amplified using the TempliPhi Amplification Kit (GE Healthcare, Chicago IL), which contains phi29 DNA polymerase (Dean et al., 2002), according to the manufacturer’s instruction at 30°C for 18 h. The reaction buffer solution of the TempliPhi Amplification Kit included random hexamer primers. Separately, 2 µL of total DNA was denatured at 95°C for 3 min, then cooled on ice for 3 min for amplification with the EquiPhi29 kit (Thermo Fisher Scientific, Waltham MA). The cooled reaction was mixed with 0.5 μL of 10X EquiPhi29 Reaction Buffer, 1.0 μL of Exo-resistant random primers, and 1.5 μL of nuclease-free water. The denatured DNA product (5 μL) was amplified using 1 μL (10 U) of EquiPhi29 DNA polymerase, 1.5 μL of 10X EquiPhi29 Reaction Buffer, 0.2 μL of 100 mM DTT, 2 μL of 10 mM dNTP mix, 1.0 μL (0.1 U) of pyrophosphatase and 9.3 μL of nuclease-free water (Povilaitis et al., 2016) according to the manufacturer’s instructions, except that the reactions were performed at 40°C for 2 h. EquiPhi29 conditions were optimized to retain the highest amount of dsDNA based on Povilaitis et al., 2016, the manufacturer’s report, and an optimization experiment (Povilaitis et al., 2016) (Supp. 1A).

Total DNA (100 ng) and RCA products (100 ng) generated using the TempliPhi Amplification Kit were treated with 1 μL (1 U) of Mung Bean nuclease (New England Biolabs, Ipswich, MA), 3 μL of CutSmart Buffer (New England Biolabs) in a 30 μL reaction volume for 30 min at 30°C. The solution was inactivated with 3 μL of SDS (0.01%), and DNA was recovered by ethanol precipitation, according to the manufacturer’s specifications. RCA products (11 μL) generated using the TempliPhi Amplification Kit were also treated with 1.0 μL (1 U) of Klenow (large subunit) (New England Biolabs) and 1.0 μL (1 U) of T4 DNA polymerase (New England Biolabs) in a mixture of 5.0 μL 10X NEB Buffer #2 (New England Biolabs), 0.5 μL 10 mM dNTP, and 31.5 μL nuclease-free water at 25°C for 1 h. The reaction was inactivated by incubation at 75°C for 1 h. After the repair reaction, residual salt and enzyme were removed by suspending 60 μL of SPRIselect beads (Beckman Coulter, Pasadena CA) in the 50 μL RCA/repair reaction. The mixture was incubated at room temperature for 5 min, placed on a DynaMag™-2 magnetic rack (Thermo Fisher Scientific, Waltham MA) for 5 min at room temperature or until the liquid was clear. The liquid was removed and the bead pellet was washed in 200 μL of 80% ethanol on the magnetic rack for 30 sec. The ethanol was removed and air-dried for 5 min on the magnetic rack. The pellet was resuspended in 17 μL of nuclease-free water.

The concentrations of the RCA products after the various treatments described above were measured using a Qubit 3.0 fluorometer with the dsDNA HS assay kit or ssDNA assay kit, according to manufacturer’s instructions (Thermo Fisher Scientific, Waltham MA). The dsDNA HS assay kit only detects dsDNA, while the ssDNA assay kit detects both ssDNA and dsDNA (i.e. total DNA). For all dsDNA concentration measurements (e.g., Figure 2), Qubit fluorometer readings were used to calculate the amount of DNA. The amount of total dsDNA (in nanograms) was calculated directly using the high sensitivity dsDNA buffer to measure the concentration (ng/μL) and multiplying by the volume of the RCA reaction. Total ssDNA mass was estimated by multiplying the concentration (ng/μL) determined using ssDNA buffer by the volume of the RCA reaction, and subtracting the total mass of dsDNA determined using high sensitivity dsDNA HS assay kit.

**Fig 2.**
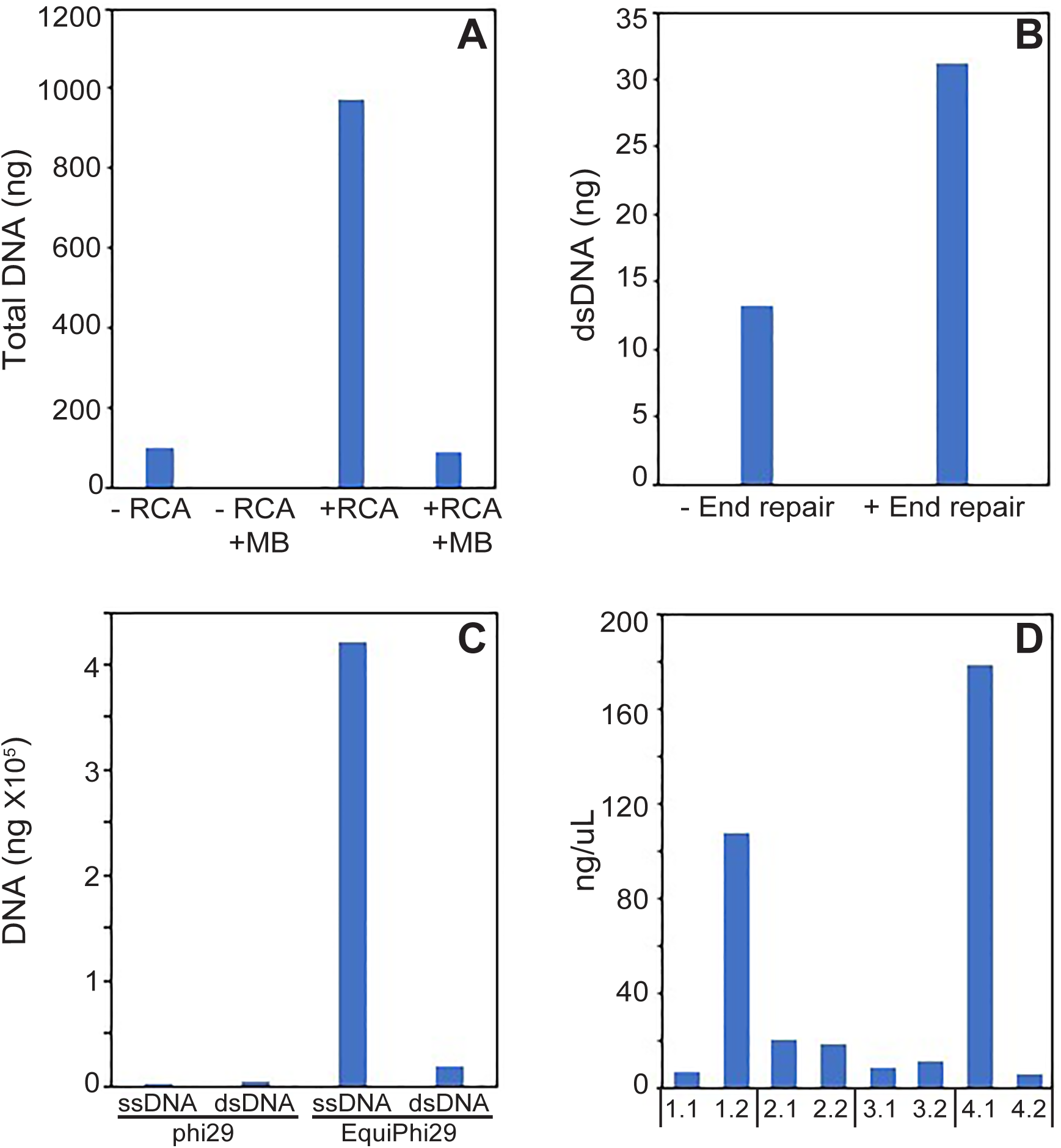
Improvements to RCA reactions. (A) The amount of total DNA (-RCA) and total DNA amplified with RCA (+RCA) treated (+MB) or not treated (-MB) with Mung Bean nuclease (MB). (B) The amount of DNA (ng) in an RCA reaction before -end repair and after +end repair. (C) Amount of ssDNA and dsDNA after RCA amplification of total DNA with either Phi29 or EquiPhi29 DNA polymerases. (D) The concentration of the total DNA of technical replicates from 4 cassava leaf DNA samples amplified using RCA.

### 2.5 Library Preparation

Total DNA, size-selected DNA, or virion DNA (2 μL) from Kibandameno leaves, tomato leaves, or whiteflies was amplified using EquiPhi29 DNA polymerase as described above. Two separate reactions were set up for each sample. Each RCA reaction was treated with Klenow and T4 DNA polymerases followed by a purification step using SPRIselect beads, as described above. After the clean-up step, each RCA reaction was diluted to 0.2 ng/μL (1 ng total in 5 μL) for library construction. Libraries were prepared using the Nextera XT DNA Library Prep Kit (Illumina, San Diego CA) following manufacturer’s instructions using Unique Dual Index adaptors (Integrated DNA Technologies, San Jose CA) and 12 rounds of PCR. The Nextera XT DNA Library Prep Kit was chosen because of its rapid library preparation, low input of DNA, and optimization for small genomes. The libraries were cleaned with 40 μL of SPRIselect beads, as described above. The libraries were analyzed for size distribution (400-800 bp), yield, and quality using a 2100 Bioanalyzer Instrument (Agilent Technologies, Santa Clara, CA). The concentration of each library was determined using a Qubit 3.0 fluorometer using the dsDNA HS assay kit as described above. Libraries were diluted to 15 nM, and equal molar amounts were pooled for sequencing on an Illumina MiSeq platform.

The molarity of each library was determined using the following formula: (ng/μL)/(nmol/μL) x (1 x 10^6^). The molecular weight of each library was determined by taking the average base-pair length from the Bioanalyzer profile multiplied by the average mass of the four nucleotide bases plus the weight of 5’-PO4 ((average base-pair length x 607.8) + 157.9).

The average mass of nucleotide bases and the MW of 5’-PO4 was taken from Thermo Fisher Scientific DNA and RNA Molecular Weight and Conversion guide (Scientific). Libraries were pooled for sequencing on an Illumina MiSeq instrument (Fig. 1). A full printable version of our protocol is available at cassavavirusevolution.vcl.ncsu.edu.

### 2.6 Data Analysis

Raw sequencing data were processed using Cutadapt (v.1.16.3) to remove the universal 3’ adapters from the paired-end reads and to trim the 5’ ends to give fastq quality scores > 30 (Martin, 2011). The quality-controlled reads were aligned to the DNA-A and DNA-B components of ACMV and EACMCV or ToMoV using BWA mem (v. 0.8) with default parameters (Li, 2013; Li et al., 2009). Quality-controlled reads were also aligned to the cassava reference genome v.7 (Bredeson et al., 2016) for cassava plant samples, to the tomato reference SL4.0 assembly (Hosmani et al., 2019) for tomato samples, and to MEAM1 assembly (GenBank ASM185493v1) (Chen et al., 2016) for whitefly samples. Duplicate reads were discarded using Picard MarkDuplicates (v. 2.18.2.1, http://broadinstitute.github.io/picard/). Samtools idxstats (v.2.0.3) was used to generate mapping statistics (Li et al., 2009). Sufficient read coverage was designated as 1000X fold coverage (∼20,000 reads/genome) based on (Juárez et al., 2019). Coverage was calculated using the following formula from Illumina (coverage=read length X number of reads/genome length). For workflow and full parameters see Galaxy workflow, ViralSeq (cassavavirusevolution.vcl.ncsu.edu) (Giardine et al., 2005). Raw Illumina data are available at the NCBI Sequence Read Archive (PRJNA658475).

## 3. Results

### 3.1 Analysis of RCA variability

Begomoviruses have circular ssDNA genomes that are converted to dsDNA during viral replication in plants. Viral ssDNA accumulates to high levels during infection, while viral dsDNA occurs at much lower levels. Many library protocols for short-read sequencing involve the ligation of adapters to dsDNA ends or transposase-mediated fragmentation and tagging of dsDNA. Thus, the *in vitro* conversion of viral ssDNA to dsDNA is the first step during library construction for ssDNA viruses. RCA is used frequently to amplify circular viral DNA genomes and to convert viral ssDNA to dsDNA (Inoue-Nagata et al., 2004; Dean et al., 2001). We examined the efficiency of RCA to convert ssDNA to dsDNA and several parameters that might influence the amount of virus-specific dsDNA available for library construction.

Total DNA isolated at 28 dpi from four symptomatic Kibaha plants infected with ACMV and EACMCV was incubated in RCA reactions containing phi29 and random hexamers (TempliPhi Amplification Kit). The RCA reactions and an equal amount of unamplified total DNA were digested with Mung Bean nuclease (MB) to remove ssDNA from the samples. RCA increased the amount of total DNA 10-fold relative to input (Fig. 2A). The amounts of the input DNA and the DNA after RCA were both greatly reduced by MB treatment, indicating that most of the DNA before and after RCA is single-stranded and therefore cannot be ligated to library adapters (Fig. 2A).

For successful library construction, we sought to increase the amount of viral dsDNA after RCA. During the RCA de-branching step, ssDNA overhangs may be present on dsDNA products, preventing adaptor ligation and leading to exclusion from the final library. To decrease ssDNA overhangs after RCA, we used an end repair reaction to convert ssDNA overhangs to dsDNA. Using the same total DNA sample in Fig. 2A, RCA was performed followed by an end-repair by T4 polymerase and DNA polymerase I (Klenow fragment). The end-repair reaction increased the amount of dsDNA 2-fold (Fig. 2B), indicating that repairing the debranched ends increased the amount of dsDNA. We also tested random primers versus virus-specific primers in the RCA reaction and found that random primers resulted in equal or higher levels of dsDNA depending on the concentration of virus-specific primers, supporting findings by Dean *et al*., 2001 (Supp. 1B).

To further increase the amount of dsDNA after RCA, we tested a modified form of phi29 DNA polymerase marketed as EquiPhi29, which has been reported to increase dsDNA output up to 7-fold (Povilaitis et al., 2016). Total DNA was amplified in RCA reactions containing the phi29 or the EquiPhi29 DNA polymerase. The amounts of dsDNA and ssDNA were measured using the Qubit dsDNA HS assay kit and ssDNA assay kit as described in the methods. The EquiPhi29 DNA polymerase yielded ∼175-fold more ssDNA and 5-fold more dsDNA than the phi29 DNA polymerase (Fig. 2C). Even though most of the increase in RCA products was ssDNA, the 5-fold increase in dsDNA when combined with the 2-fold increase after the DNA end-repair reaction resulted in more dsDNA available for ligation to library adapters.

During the process of testing different parameters, we noticed that RCA was highly variable in DNA output. Four total DNA samples (1-4) isolated from symptomatic Kibandameno leaves were amplified using RCA with EquiPhi29 and phi29 (Fig. 2D). This process was repeated twice generating two technical replicates for each sample. The dsDNA concentration of each sample and its technical replicate were measured using a Qubit dsDNA HS assay kit (Fig. 2D). Two of the four samples had technical replicates that differed in concentration by more than 10-fold (samples 1 and 4, Fig. 2D). Variability was also observed with phi29 (data not shown). This problem was overcome by standardizing the amount the RCA product (2 μL at 5 ng/μL, i.e. 10 ng DNA) used for library construction. If the yield of the RCA product from a given reaction was insufficient for dilution to 5 ng/μL, that reaction was repeated.

### 3.2 Total DNA versus size-selected DNA from leaf tissue

Our goal was to develop a protocol where we could achieve sufficient read coverage to detect viral variants across a complete transmission cycle. Based on the coverage reported by Juárez et al., (2019) to detect viral variants and our calculations, we set 1000X coverage (ca. 20,000 150-bp reads/genome component) as our minimum coverage for detecting low viral variants occurring at 3% and 1% frequency in the population (for 30 and 10 variant-supporting reads, respectively). To achieve this goal, we examined additional methods to increase the number of reads mapping to the viral genomes.

The DNA-A and DNA-B components of the ACMV and EACMCV genomes are ca. 2.8 Kb in size and, as such, are much smaller than plant genomic DNA. Hence, we asked if size selection would increase the number of reads mapping to the viral genomes. We used the BluePippin system to select for DNA < 6 Kb from total DNA samples isolated from symptomatic Kibandameno leaves from two plants. The starting total DNA and the size-selected DNA were amplified using the optimized RCA protocol. Libraries were generated using our experimental pipeline (Fig. 1) and sequenced on the Illumina MiSeq platform. We sequenced two technical replicates for each sample. After processing and mapping the resulting reads to the ACMV and EACMCV reference genomes and the cassava reference, we found that nearly all of the reads mapped to the viral reference genomes when the DNA was size-selected (blue) compared to half of the reads for total DNA (light grey) (Fig. 3A). Generally, the use of size-selected DNA (blue) for library construction increased the number of reads mapping to each of the viral genome components compared to total DNA (grey), improving mapping by ∼ 2-fold (Fig. 3B). Nearly all of the reads that did not map to the viral genome components but mapped to the cassava reference genome (Fig. 3A), indicating that size selection is an effective method for separating viral DNA from host DNA. Size-selection was effective in increasing the viral read counts in samples with both high and low levels of CMB genome components (Supp. 2A and B), indicating that size selection can produce reliable results over a 10-fold range. However, the virus titer before RCA does not reflect the resulting coverage and read count after RCA and NGS sequencing (compare Supp. 2A and 3). This result also underscores the variability of RCA (Fig. 3C).

**Fig 3.**
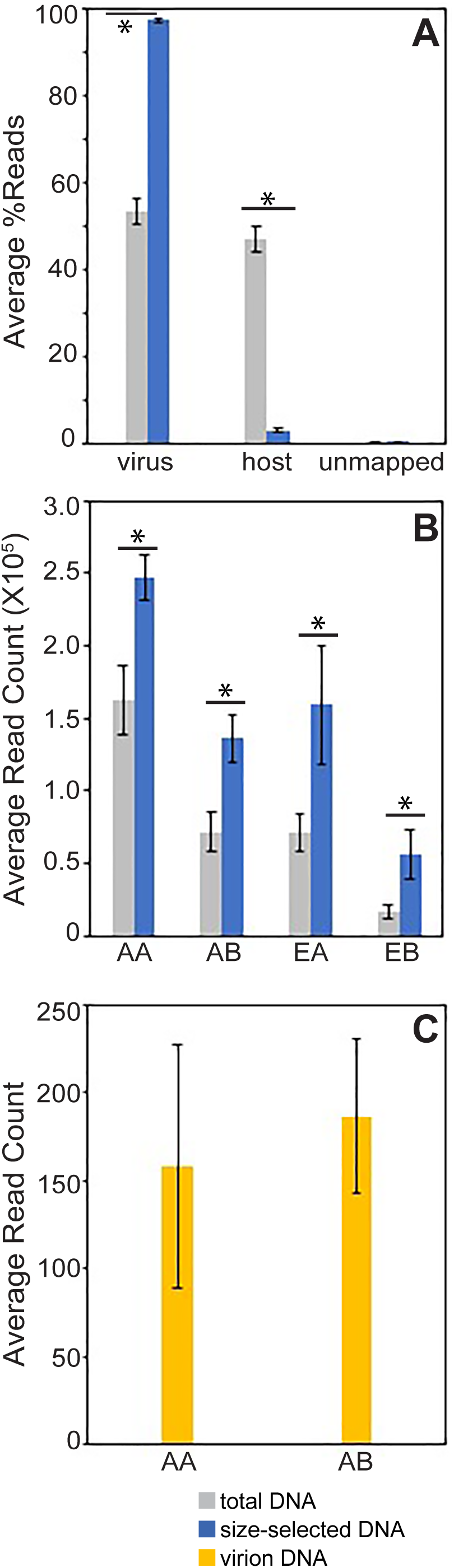
Size selection increases viral DNA read counts for cassava samples. (A) The average percent reads mapping to viral DNA (ACMV and EACMCV) and host DNA, and unmapped reads for total DNA (grey) and size-selected DNA (blue) samples. (B) The average number of reads corresponding to ACMV DNA-A (Hosmani et al.), ACMV DNA-B (AB), EACMCV DNA-A (EA), and EACMCV DNA-B for total (grey) and size-selected (blue) DNA samples. (C) The average number of reads corresponding to ACMV DNA-A (Hosmani et al.), ACMV DNA-B (AB) for virion DNA (yellow). The bars in A, B correspond to 2 standard errors from two bio-samples and two technical reps each. The bars in C correspond to 2 standard errors from three bio-samples with two technical replicates each. The asterisks indicate a significant difference (P-value < 0.05) between the viral read counts in size-selected and total DNA in Student’s t-tests.

Size-selection also improved read coverage compared to coverage from total DNA for both cassava (Supp. 3A-D) and whitefly (Supp. 3E-H) samples. The coverage was relatively even across the genomes. Dips in coverage were seen in some profiles at the 3’ ends of the convergent transcription units (at the ends of the converging arrows) and in the 5’ intergenic regions (at the ends of the linear maps), but the coverage was still above 1000X.

After observing that size selection increased viral read count, we evaluated whether virion DNA containing only packaged viral ssDNA would also increase viral read count (Fig. 3C). The resulting average read count (yellow) from three bioreplicates was highly variable and 500-1000 fold lower than the average read counts of total or size-selected DNA (Fig. 3B).

### 3.3 Sequencing viral DNA from viruliferous whiteflies

We used the ToMoV-tomato-whitefly pathosystem to assess if our optimized protocols could be applied to another begomovirus. As with the cassava pathosystem, sequencing libraries generated from total DNA (grey) isolated from a ToMoV-infected tomato plant (source plant) resulted in fewer viral reads than libraries constructed from size-selected DNA (blue; Fig. 4A). The average reads from virions for ToMoV DNA-A were similar to total DNA and less than size-selected DNA, while the average read counts from virions for ToMoV DNA-B were lower than total and size-selected DNA (Fig. 4A). This underscores the variability of sequencing from virions. Examining the average percent reads mapping to ToMoV versus host DNA (source plant), we found that over 99% of the reads mapped to ToMoV when using size-selected DNA (Fig. 4B). Using total DNA and virion DNA resulted in ∼30% and ∼25% of the reads mapping to ToMoV, respectively. This confirms the advantage of size-selection seen in the cassava pathosystem (Fig. 3).

**Fig 4.**
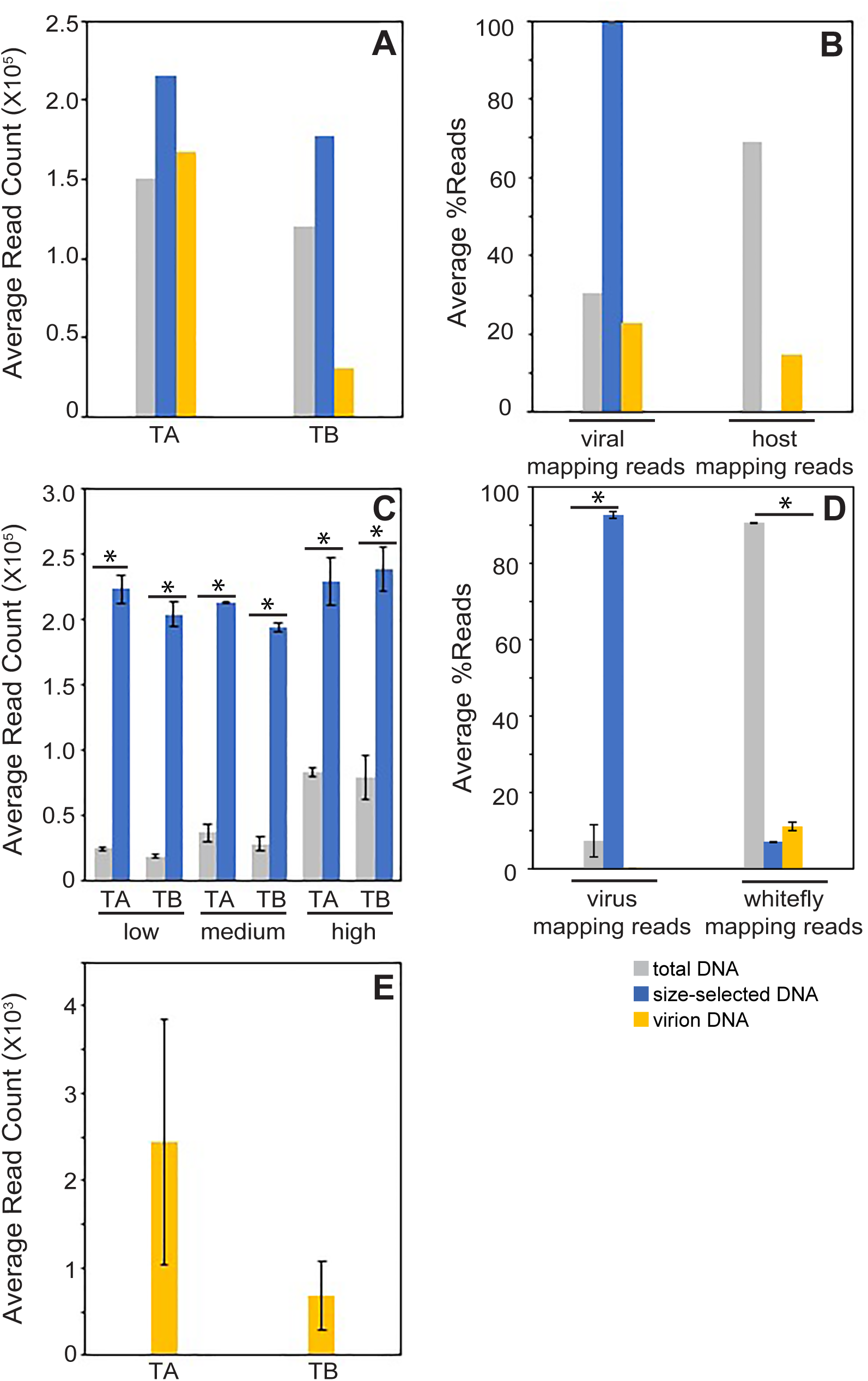
Size selection increases viral DNA read counts for tomato and whitefly samples. A) The average number of reads corresponding to ToMoV DNA-A (TA) and ToMoV DNA-B (TB) for total (grey), size selected (blue), virion (yellow) DNA samples from the tomato source plant. (B) The average percent reads mapping to viral DNA (ToMoV) and host DNA for total DNA (grey), size-selected DNA (blue), and virion DNA (yellow) samples. (C) The average number of reads corresponding to ToMoV DNA-A (TA) and ToMoV DNA-B (TB) for total (grey) and size-selected (blue) DNA samples from pools of 5 whiteflies with low (30,000-40,000 DNA-A copies/ng total DNA), medium (45,000-60,000 DNA-A copies/ng total DNA), and high viral loads (150,000-200,000 DNA-A copies/ng total DNA) virus loads. (D) The average percent reads mapping to viral DNA (ToMoV), host DNA, and whitefly DNA for total DNA (grey), size-selected DNA (blue), and virion DNA (yellow) samples. (E) The average number of reads corresponding to ToMoV DNA-A (TA) and ToMoV DNA-B (TB) for virion DNA (yellow). The bars in A and C correspond to 2 standard errors for three bio-samples with two technical replicates. The asterisks indicate a significant difference (P-value < 0.05) between the viral read counts in size-selected and total DNA in Student’s t-tests.

We also sequenced DNA from groups of 5 whiteflies that had acquired ToMoV virions from the sampled source plant. We sequenced groups of whiteflies with low, medium, and high viral loads as determined by qPCR. The low, medium and high load groups were split into total DNA and size-selected DNA treatments and sequenced. Read counts for the total DNA samples increased with viral load (Fig. 4C) and were lower than read counts for total DNA from infected tomato (Fig. 4A). In contrast, read counts for size-selected DNA samples (blue) were similar across the different viral load levels and were 20-fold higher than the total whitefly DNA samples (grey; Fig. 4B). Overall, the percent of reads mapping to the viral genome was greater than 90% for size-selected DNA (Fig. 4D). Size-selection also resulted in a 6-fold increase in read coverage compared to total DNA (Supp 3E-H).

We also attempted to sequence virion DNA from ToMoV-infected tomato (Fig. 4A) and three sets of five viruliferous whiteflies (Fig. 4E). The resulting reads for virion DNA (yellow) from the ToMoV-infected tomato were variable between the DNA-A and DNA-B components, and the average read count was 100-fold lower than total DNA (grey) and size-selected DNA (blue). The resulting reads from virion DNA (yellow) from viruliferous whiteflies were less than 1000X coverage and varied between bioreplicates, indicating that sequencing from whitefly virions was not reproducible. It is also important to note that sequencing total and size-selected DNA provides information both viral ssDNA and dsDNA while sequencing virion DNA only yields sequence data about ssDNA.

## 4. Discussion

In this study, we examined the impact of RCA reaction conditions and size selection on short-read sequencing of begomovirus DNA. Our optimized protocol effectively enriched for viral DNA and produced short-read sequence data from enhanced RCA reaction products across a range of viral loads. Using size-selected DNA, over 90% of the reads mapped to the viral reference genomes from cassava (Fig. 3A). Without size selection, ca. 50% of the reads mapped to the viral genome and the remaining 50% mapped to the host genome (Fig. 3A). Given that both approaches can yield high numbers of viral reads, using total DNA for library construction may be the preferred approach for plant samples when access to size fractionation instrumentation, cost, and/or time are limiting. A recent study similarly concluded that size selection can be beneficial for long-read sequencing of begomoviruses, which has advantages for certain applications (Mehta et al. 2019).

Previous studies using RCA and short-read sequencing to characterize begomovirus sequences reported less than 50% of the reads mapping to the viral genome and even lower proportions when the RCA step was omitted (Kathurima, 2016). Sequencing of CMBs yielded mapping ranges of 0.87-6.9% from cassava (Kathurima, 2016) and 0.9-29.6% from *Nicotiana benthamiana* (Chen et al., 2019). Low read counts following RCA were also observed for tomato leaf curl New Delhi virus (0.47-1.05 %; (Juárez et al., 2019) and euphorbia yellow mosaic virus (1.24-1.35%; (Richter et al., 2016). Our improved sequence method resulted in at least 7X higher viral mapping reads for CMBs and 66X higher than other begomoviruses reported in the literature.

Our methods can be used to characterize viral DNA sequences in whiteflies (Fig. 4B). Size selection had a much larger impact on viral read counts from viruliferous whitefly samples compared to infected plant samples, resulting in at least 1000X coverage across a range of viral loads. Sequencing total DNA from whiteflies results in 1000X read coverage when viral loads are high but not when they are low. In contrast, a wide range of viral loads was successfully sequenced using both size selection and total DNA approaches from infected plants. Sequencing viral sequences from whitefly virions produced average viral read counts that were highly variable and 100-fold lower than that average read count from total and size-selected DNA (Fig. 4C and E). Currently, most NGS sequencing from whiteflies is from enriched virions for vector-enabled metagenomic surveys (Ng et al., 2011; Rosario et al., 2015). We found that sequencing from virions did not provide enough coverage and read depth for studying virus population diversion in pools of 5 whiteflies.

We found that RCA was not efficient at converting ssDNA to dsDNA (Fig. 2A). Even with an improved DNA polymerase (EquiPhi29), most of the RCA product is ssDNA. RCA has a preference to amplify ssDNA as linear, concatenated copies with low conversion of ssDNA to dsDNA, which can be improved by increasing the reaction incubation time to over 25 hours (Ducani, Bernardinelli, and Högberg, 2014; Zhang and Tanner, 2017). However, the Equiphi29 DNA polymerase resulted in ∼5-fold more dsDNA after RCA with only a 2-hour incubation time, increasing the amount of viral DNA template available for library construction in much shorter reaction time (Fig. 2C). We also found that the amount of DNA produced by RCA varied even when reactions contained the same input DNA (Fig. 2D). Thus, it is important to standardize the amount of RCA product used for library preparation.

## 6. Conclusion

We established a short-read sequencing protocol for ssDNA viruses that provides high numbers of reads that map to viral reference genomes. The method can use total DNA or size-selected DNA from leaf and whitefly samples that are amplified using random primers and a modified phi29 DNA polymerase prior to library construction. We also found that RCA is variable and is strongly biased towards the amplification of ssDNA products. We cannot fully explain the poor conversion rate to dsDNA, and further studies are needed to understand how ssDNA is converted to dsDNA during RCA. In summary, we have developed an improved tool for begomovirus DNA diagnostics and studying begomovirus population dynamics. Our approach should be applicable to other CRESS DNA viruses.

## Author contributions

CDA - Experimental design, execution, and analysis; manuscript preparation

JSH - Experimental design and data analysis; manuscript preparation

AED - Experimental design, execution, and data analysis; manuscript preparation

DOD - Experimental execution

IC - Data analysis

SD-Experimental design

LHB - Experimental design and manuscript preparation

## Acknowledgments

We thank Mary Beth Dallas for her help growing cassava and tomato plants. The Office of Advanced Research Computing (OARC) at Rutgers, The State University of New Jersey, provided access to and maintenance of the Amarel cluster. We also thank the NC State University Genome Science Laboratory for their support.

## Figure Legends

*Supp 1. Optimization of RCA conditions*

(A) Amount of ssDNA and dsDNA after RCA amplification of total DNA with EquiPhi29 polymerase for 2 h and 3 h at 40°C. (B) Amount of dsDNA after RCA amplified with random hexamers (hx) alone or combined with different amounts of virus-specific primers (vs).

*Supp 2. Virus levels and virus-mapping read count for cassava biological replicates*.

(A) Log viral copy number of ACMV-A (AA0, ACMV-B (AB), EACMCV-A (EA), and EACMCV-B (Bernardo et al.) for biological replicate 1 (green) and biological replicate 2 (blue).

(B) Average read count of two technical replicates for biological replicate 1 (green) and 2 (blue) for ACMV-A (Hosmani et al.), ACMV-B (AB), EACMCV-A (EA), and EACMCV-B (Bernardo et al.) without size-selection. (C) Average read count of two technical replicates for biological replicate 1 (green) and 2 (blue) for ACMV-A (Hosmani et al.), ACMV-B (AB), EACMCV-A (EA), and EACMCV-B (Bernardo et al.) with size-selection.

*Supp 3. Plots of Illumina read depth across virus segments*.

Each row corresponds to one library, from leaf tissue from a cassava plant (A to D) or a single pool of whiteflies (E to H). Pairs of rows correspond to technical duplicate libraries made from total DNA (A and B, E and F) or to size-selected DNA (C and D, G and H). Canonical virus genes are drawn as gray arrows below each set of graphs, left to right for virus sense (AV1, AV2 [for ACMV and EACMCV], BV1), and right to left for complementary sense (AC1 to AC4, BC1).

