## Supplementary figures and images for "An improved experimental pipeline for preparing circular ssDNA viruses for next-generation sequencing"

### Supplemental Figure 1

Supp 1 Aimone et al

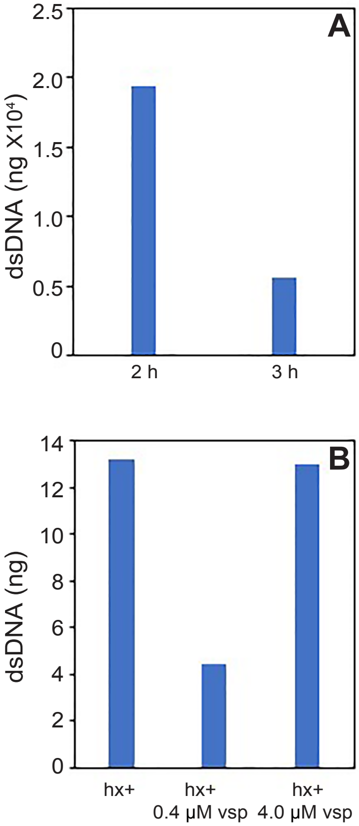

### Supplemental Figure 2

Supp 2 Aimone et al

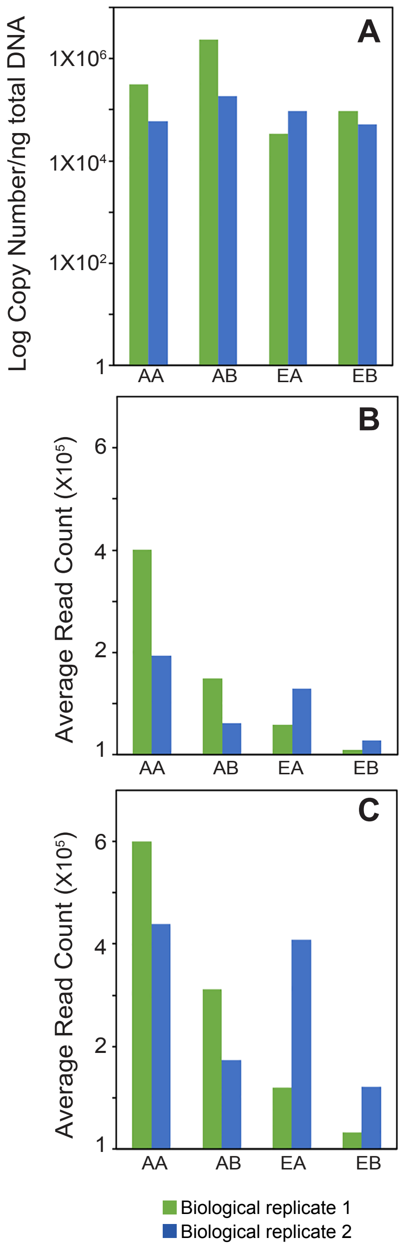

### Supplemental Figure 3

Supp 3 Aimone et al

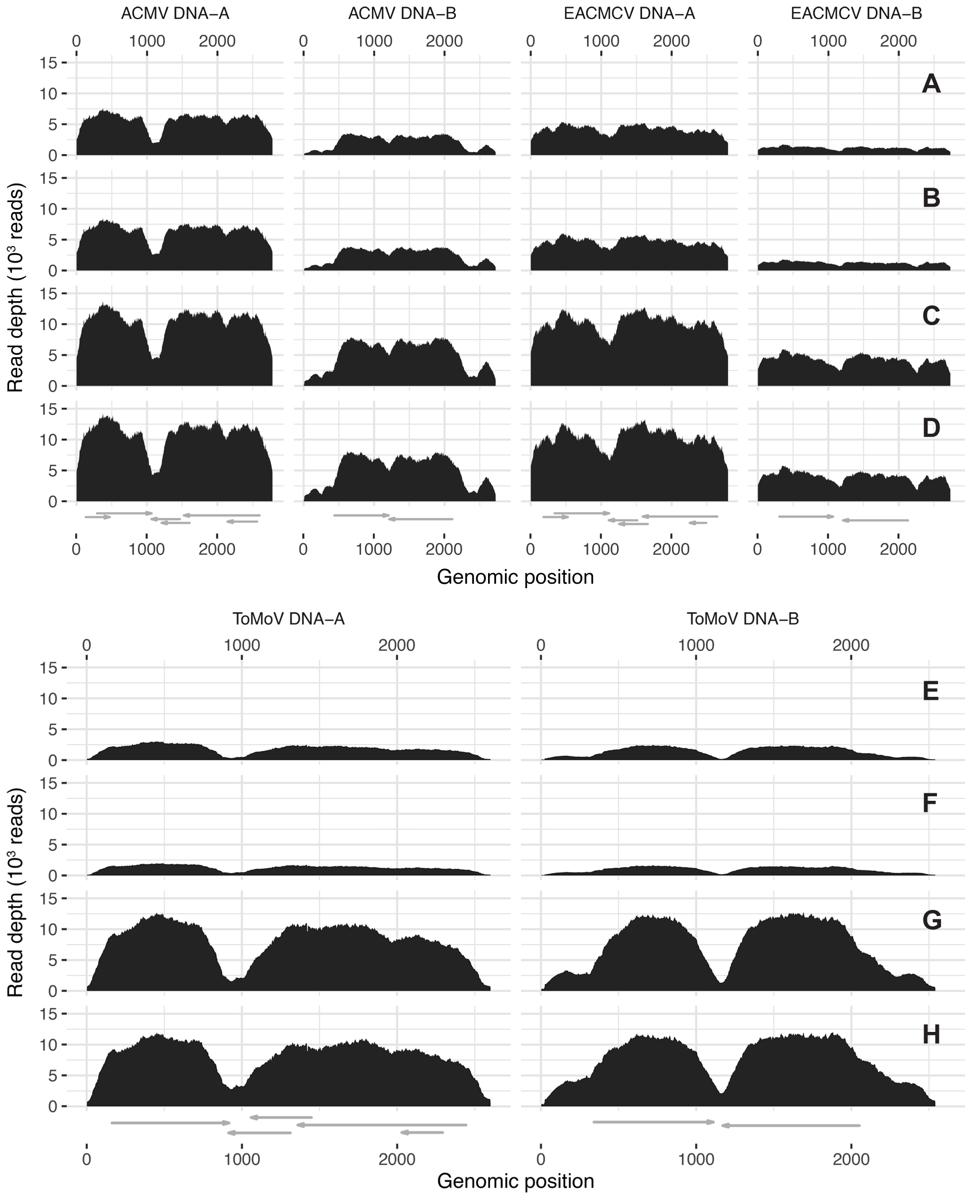
